# Revisiting the phylogeny of *Ficus* (Moraceae): When Next Generation Sequencing corroborates Past Generation Botanists

**DOI:** 10.1101/340463

**Authors:** Jean-Yves Rasplus, Lillian Jennifer Rodriguez, Christine Tollon-Cordet, Astrid Cruaud

## Abstract

Despite their ecological and evolutionary importance, fig trees still lack a resolved phylogeny. We used multiple analytical approaches and 600 conserved RAD-seq loci collected across 40 representative species to infer phylogenetic relationships among the major clades of *Ficus*. Our results converged on a fully resolved phylogeny that contradicts previous hypotheses by not placing section *Pharmacosycea* sister to all other fig trees. Instead, we find that subgenus *Sycomorus* is sister to two clades, one comprising all other gynodioecious sections, and the other containing all monoecious sections. Our results corroborate earlier botanists’ hypotheses and provide a baseline for future classification. We infer the evolution of key traits (growth form, breeding system and pollination mode) and find that, contrary to previous hypotheses, the ancestor of extant fig trees was likely a gynodioecious and actively pollinated tree. Our results open new opportunities to explore diversification patterns in fig trees and better understand their mutualistic interaction with wasps.

## 1. INTRODUCTION

*Ficus* (Moraceae) is a pantropical and hyperdiverse genus (*ca* 850 species [1]) with a large range of growth forms (trees, hemiepiphytes, shrubs, climbers) and an enormous diversity of ecologies. As figs are important food source for hundreds of frugivorous species [2], fig trees are key components of tropical ecosystems. They are also known for their intricate relationships with their pollinating wasps (Agaonidae). Indeed, since *ca* 75 Myr, fig trees and agaonids are obligate mutualists [3]. The wasp provides pollination services to the fig tree, while the fig tree provides a breeding site for the wasps, and none of the partners is able to reproduce without the other [4]. Fifty-two percent of *Ficus* species are monoecious, while 48% are gynodioecious. In monoecious species, figs contain staminate and pistillate flowers and produce pollen and seeds. Gynodioecious species are functionally dioecious with male and female functions segregated on separate individuals. *Ficus* species are pollinated either actively (two-thirds) or passively (one-third) [5]. In passively pollinated figs, emerging wasps are dusted with pollen before flying away, while in actively pollinated figs, wasps use their legs to collect pollen they will later deposit into receptive figs.

Despite its ecological importance, the evolutionary history of the genus remains unclear. Six subgenera and 20 sections were recognized by botanists (Table S1). Several studies have attempted to reconstruct the phylogeny of *Ficus* (Fig. 1) using Sanger sequencing of chloroplast markers [6]; external and/or internal transcribed spacers (ETS, ITS) [7, 8], a combination of nuclear markers [3, 9, 10] or next-generation sequencing of chloroplast genomes [11]. None of these studies successfully resolved the backbone of the phylogeny and there is no consensus on the relationships between major groups of *Ficus*. Plastome analysis provided improved resolution but failed to propose a consistent hypothesis because undoubtedly monophyletic subgenera based on morphology (*Sycomorus, Urostigma*) were recovered polyphyletic [11] (Fig 1). It should be pointed that with phylogenomic approaches, cases of cyto-nuclear discordance caused by hybridization or high plastome divergence are frequently reported [12], making the use of nuclear markers desirable.

**Figure 1:**
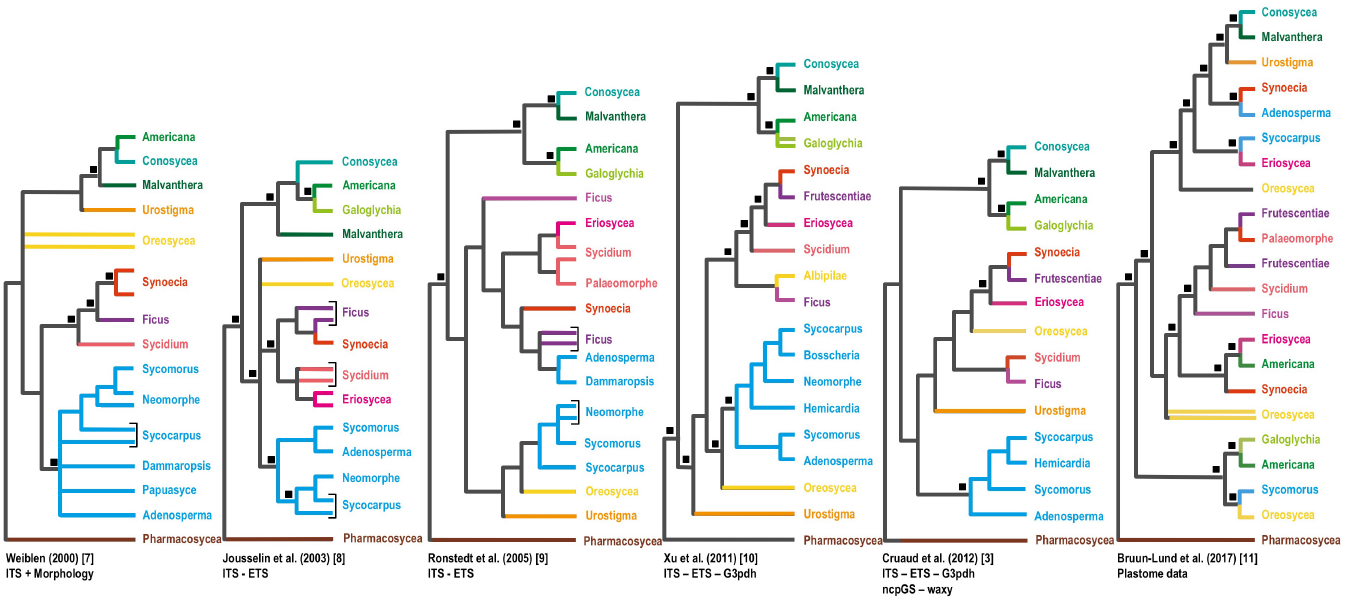
Previous hypotheses regarding the phylogeny of *Ficus*. See color legend in Fig 2.

In all studies, section *Pharmacosycea* was recovered sister to all other *Ficus*. Given the long branch leading to *Pharmacosycea* species, the phylogenetic placement of this section remains vulnerable to systematic errors. Thus, the lack of robust phylogenetic inferences and possible bias induced by systematic errors hamper our understanding of the evolutionary history of the mutualism.

Herein, we used Restriction-site-associated DNA sequencing (RAD-seq) to elucidate the phylogenetic relationships within *Ficus*. Indeed, RAD-seq has been successfully utilized to infer recent and ancient evolutionary histories of multiple organisms including plants (e.g. [13, 14]). We then use our dataset to revisit the evolution of key traits (plant habit, monoecy-gynodioecy and pollination syndromes) that may have contributed to the evolutionary and ecological success of the genus.

## 2. MATERIALS AND METHODS

Forty species of *Ficus* representing all subgenera and 17 sections as well as one outgroup were used in the study (Table S2). Library construction is detailed in Appendix S1. To sequence RAD-tags, while reducing possible LBA artefact, we used an infrequent 8-cutter restriction enzyme (*SbfI*). Cleaning of data was performed with RADIS [15] that relies on Stacks [16] for demultiplexing and removing PCR duplicates. Individual loci were built using *ustacks* [m=3; M=2; with removal (r) and deleveraging (d) algorithms enabled]. The parameter n of *cstacks* (number of mismatches allowed between sample loci when building the catalog) was set to 14 to cluster enough loci for the outgroup, while ensuring not to cluster paralogs in the ingroup. To target loci with slow or moderate substitution rate, only loci present in 75% of the samples were analysed. Loci for which samples had three or more sequences were removed. Phylogenetic analyses were performed with maximum likelihood (RAxML [17]), Bayesian (MrBayes [18]) and parsimony (TNT, [19]) approaches (concatenated, unpartitioned dataset) as well as a gene tree reconciliation method (ASTRAL-III, [20]). Stochastic mapping [21] was utilized to estimate the ancestral state and number of transitions for three traits (plant habit, monoecy-gynodioecy and pollination syndromes). Three models were tested: equal rates (ER), symmetric (SYM) and all rates different (ARD). The AIC score and Akaike weight for each model were computed. See Appendix S1 for further details.

## 3. RESULTS

The final dataset included 600 loci of 113 bp each for a total of 67,800 bp (44.2% GC, 17.7% missing data, 23.6% variable and 10.0% parsimony informative sites). Maximum likelihood (ML) and Bayesian (BI) analyses produced identical and highly supported topologies (Fig 2A, S1). The multispecies coalescent analysis (CO) produced a similar topology, though less resolved, which is not surprising given the short length of loci. In ML, BI and CO topologies, section *Pharmacosycea* was recovered nested within a clade of monoecious *Ficus* with high support in ML and BI. *Ficus* was subdivided in three clades: 1) subgenus *Sycomorus*, 2) the rest of the gynodioecious *Ficus* and 3) all other monoecious *Ficus*. On the contrary, in parsimony analyses, section *Pharmacosycea* was sister to all other *Ficus* (Fig S1), *Sycomorus* was nested in a clade with other gynodioecious species, while *Oreosycea* (monoecious) was sister of this gynodioecious clade. In all analyses, all subgenera, except *Ficus* and *Pharmacosycea*, were recovered monophyletic. Notably, *Ficus carica* (subgenus *Ficus*) was sister to *Sycidium* and a strongly supported clade containing *Synoecia, Eriosycea* and *Frutescentiae* was recovered. For the first time, subgenus *Urostigma* appeared monophyletic. Stochastic mapping on the Bayesian tree (Fig 2B, Table S3) revealed that the ancestor of all extant *Ficus* was likely an actively-pollinated, gynodioecious tree from which hemiepiphytes and root climbers evolved. Monoecy appeared at least twice. Active pollination was lost several times independently and never reacquired (Fig 2B).

**Figure 2:**
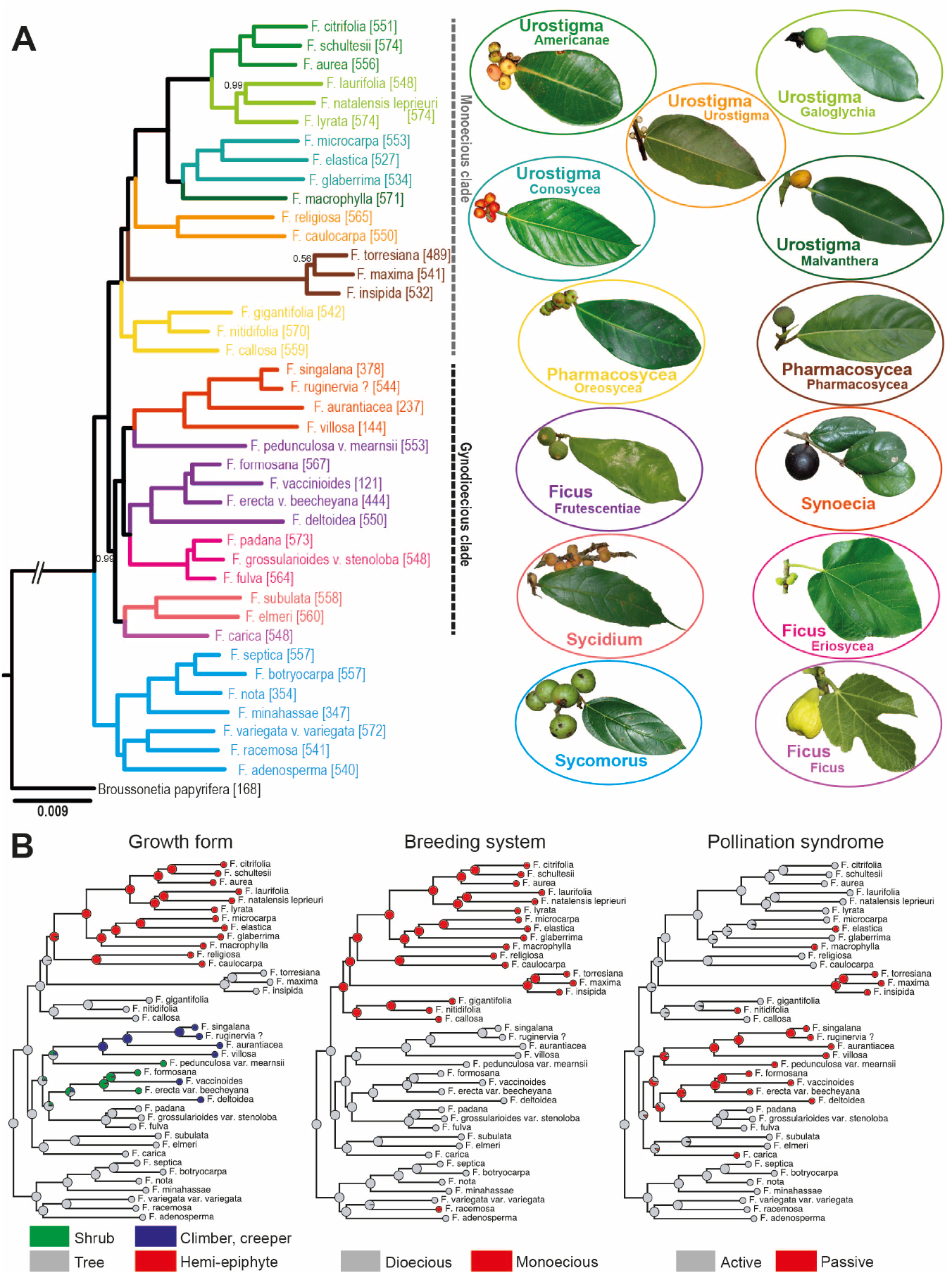
A) Bayesian topology. Posterior probability < 1.00 indicated at nodes; number of RAD-tags indicated between brackets; Subgenera / Sections illustrated with pictograms. **B) Reconstruction of traits evolution under stochastic mapping**.

## 4. DISCUSSION

Our results contradict all previous studies (Fig 1) that recovered *Pharmacosycea* sister to all other *Ficus* species. Here, only parsimony inferred this relationship (Fig S1), which suggests that early analyses were flawed by LBA. Parsimony does not model variation of evolutionary rates among sites and is thus more sensitive to LBA [22]. In fact, the relative sensitivity of parsimony and probabilistic methods to LBA is recognized as a valuable method in detecting this artefact [22]. By selecting conserved loci (8-cutter, 75%-complete matrix, ~75% similarity between individual loci for clustering), we possibly reduced the impact of higher substitution rates in section *Pharmacosycea* (a long branch is still visible, Fig S1) to a level compatible with the Gamma distribution used to model rate variation in a probabilistic framework. Flaws in the parsimony analysis raise doubt on the other statistically supported incongruences (position of *Sycomorus* and *Oreosycea*) that may have also resulted from inference bias. Therefore, we based our discussion and inferences on evolution of life history traits on the topology inferred by ML, Bayesian and Coalescence analyses.

Interestingly, this topology corroborates earlier botanists’ hypotheses suggesting that sections *Pharmacosycea* and *Oreosycea* were closely related [23], though maybe not sister taxa. Indeed, these sections have similar vegetative and floral structures such as branched stigmas and similar position of the interfloral bracts. This topology is also the first to group all monoecious fig trees (subgenera *Pharmacosycea* and *Urostigma*), a pattern also suggested by earlier botanists [24]. Relationships within the gynodioecious clade make sense morphologically, though they do not support the traditional classification. The most debatable result is the sister position of *F. carica* and *Sycidium* though it is corroborated by the small pistillode (abortive ovary in male flower) and several other characters *F. carica* shares with some *Sycidium* species [1].

Previously, monoecy was supposed to be the ancestral character-state in *Ficus*, with one (or more) change(s) to gynodioecy [7, 25, 26] and reversals to monoecy in *Sycomorus*. Passive pollination was considered the ancestral condition for the mutualism [26] and a scenario with a single shift to active pollination and several independent reversals was considered likely. However, this hypothesis relied on the fact that, in most studies, *Tetrapus* species, which are passive pollinators, were sister to all other agaonids. At best, ancestral pollination mode was considered as equivocal and one analysis (ML on *Ficus* phylogeny) favored active pollination [3]. Finally, no investigation on the ancestral growth form has been conducted. Our results shed a new light by suggesting that the ancestor of *Ficus* was most likely an actively-pollinated, gynodioecious tree. On previous phylogenies, growth forms and breeding systems were highly conserved within sections but appeared over-dispersed [7]. Our phylogeny indicates that these traits are indeed more conserved than previously thought. For example, monoecy only evolved once in non-*Sycomorus* fig trees and hemiepiphytism evolved only twice in *Urostigma* and *Sycidium*.

Clearly, this study reveals novel relationships within *Ficus*. Our first goal was to test whether RAD-tags would be conserved enough to solve relationships in an old group and reduce possible systematic bias. Increasing sampling is clearly required to ascertain our hypothesis. If our results are confirmed, the infrageneric classification will have to be reassessed. Until now, testing cospeciation between partners was hampered by the poor resolution of their phylogenies. RAD-seq may contribute in circumventing this lack of resolution. This is ideal as the understanding of the diversification of this peculiar mutualism will benefit from better-resolved trees.

## ACKNOWLEDGEMENTS

We thank A. Bain and R.A.S. Pereira for providing samples, S. Santoni for technical advices, MGX for sequencing, the Genotoul platform for computing resources and C.C. Berg and E.J.H. Corner for their eternal contributions.

## AUTHOR CONTRIBUTIONS

J.Y.R., A.C. conceived the study, analyzed data and drafted the manuscript. L.J.R., C.T.C., A.C. performed lab work. All authors gave final approval for publication.

## DATA ACCESSIBILITY

Cleaned forward reads are available as a NCBI Sequence Read Archive (#SRP149690).

## FOOT NOTE

Supplementary data are available online COMPETING INTERESTS. We declare having no competing interests.

## FUNDING

This work was partially funded by UPD-OVCRD (171715 PhDIA) and UPD-NSRI (BIO-18-1-02).

## Supporting Information to

### Revisiting the phylogeny of *Ficus* (Moraceae): When Next Generation Sequencing corroborates Past Generation Botanists

Jean-Yves Rasplus, Lillian Jennifer Rodriguez, Christine Tollon-Cordet, Astrid Cruaud.

**Appendix S1. Supplementary methods**

**Table S1:**
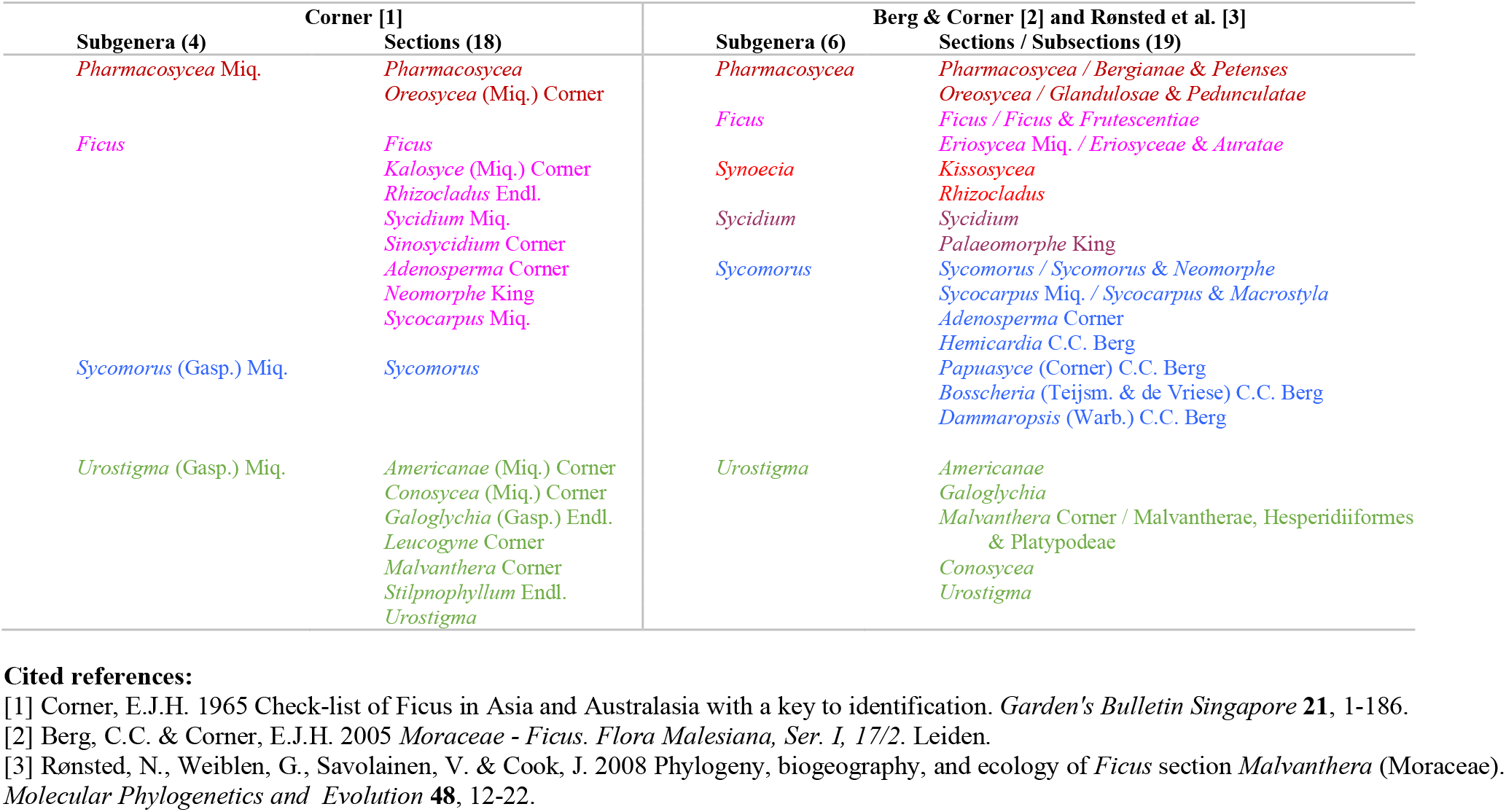
Classification of the genus *Ficus*.

**Table S2:**
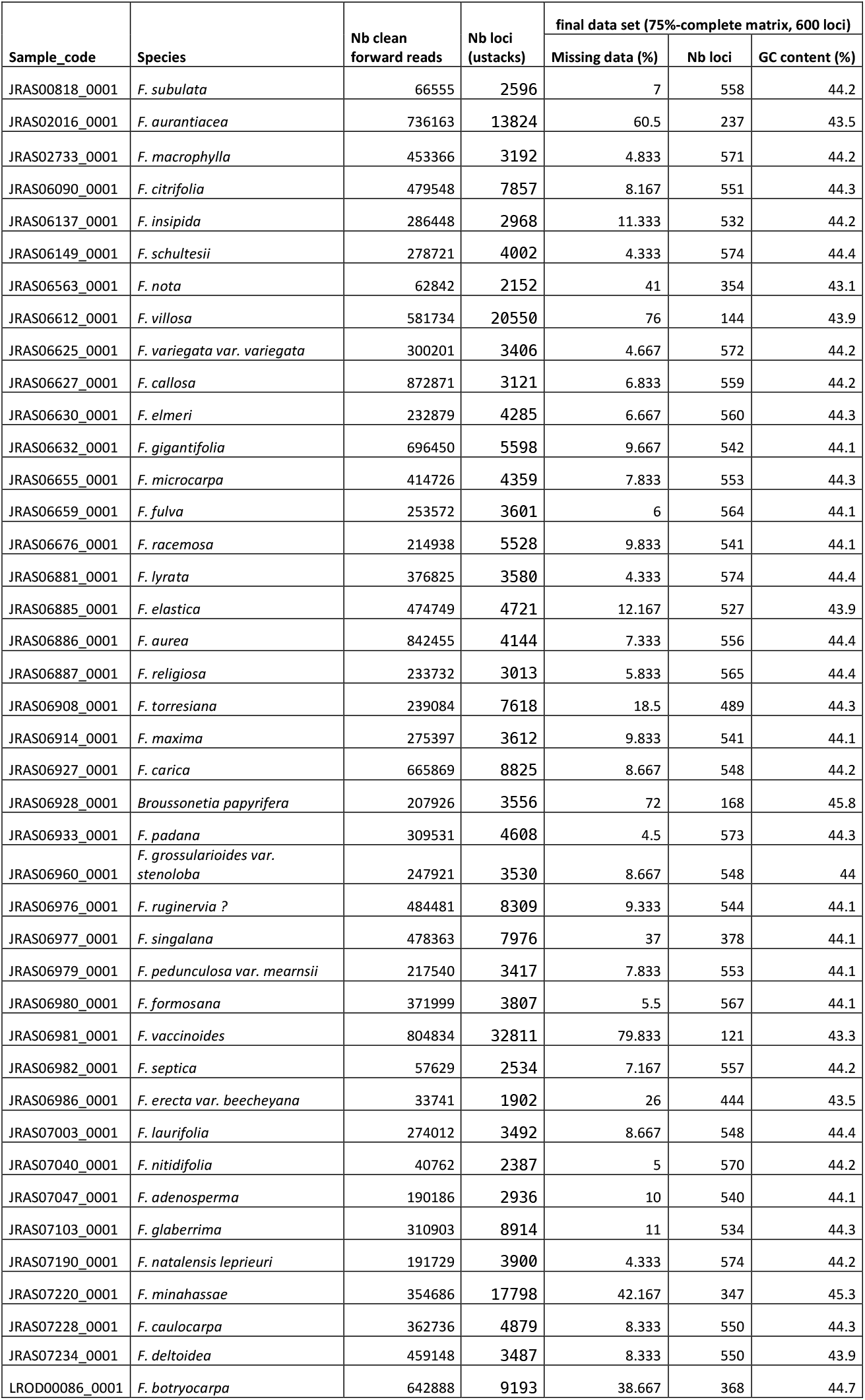
Sequencing data obtained for each sample

**Table S3:** log likelihoods, AIC scores and Akaike weights for the different models tested for stochastic character mapping. Based on these results, the following models were chosen: pollination syndrome and breeding system = ER; life form = SYM.

|  | breeding system |  |  | pollination syndrom |  |  | life form |  |  |
| --- | --- | --- | --- | --- | --- | --- | --- | --- | --- |
| model | logL | AIC | Akaike weight | logL | AIC | Akaike weight | logL | AIC | Akaike weight |
| ER | -12.24639 | 26.49278 | 0.934535666 | -26.17722 | 54.35443 | 0.986615690 | -31.63982 | 65.27964 | 0.132659624 |
| SYM | -12.94462 | 31.88924 | 0.062916983 | -29.15684 | 64.31368 | 0.006784622 | -27.76458 | 61.52916 | 0.865257204 |
| ARD | -13.15138 | 38.30277 | 0.002547351 | -26.18447 | 64.36895 | 0.006599688 | -30.79372 | 73.58743 | 0.002083171 |

**Figure S1.**
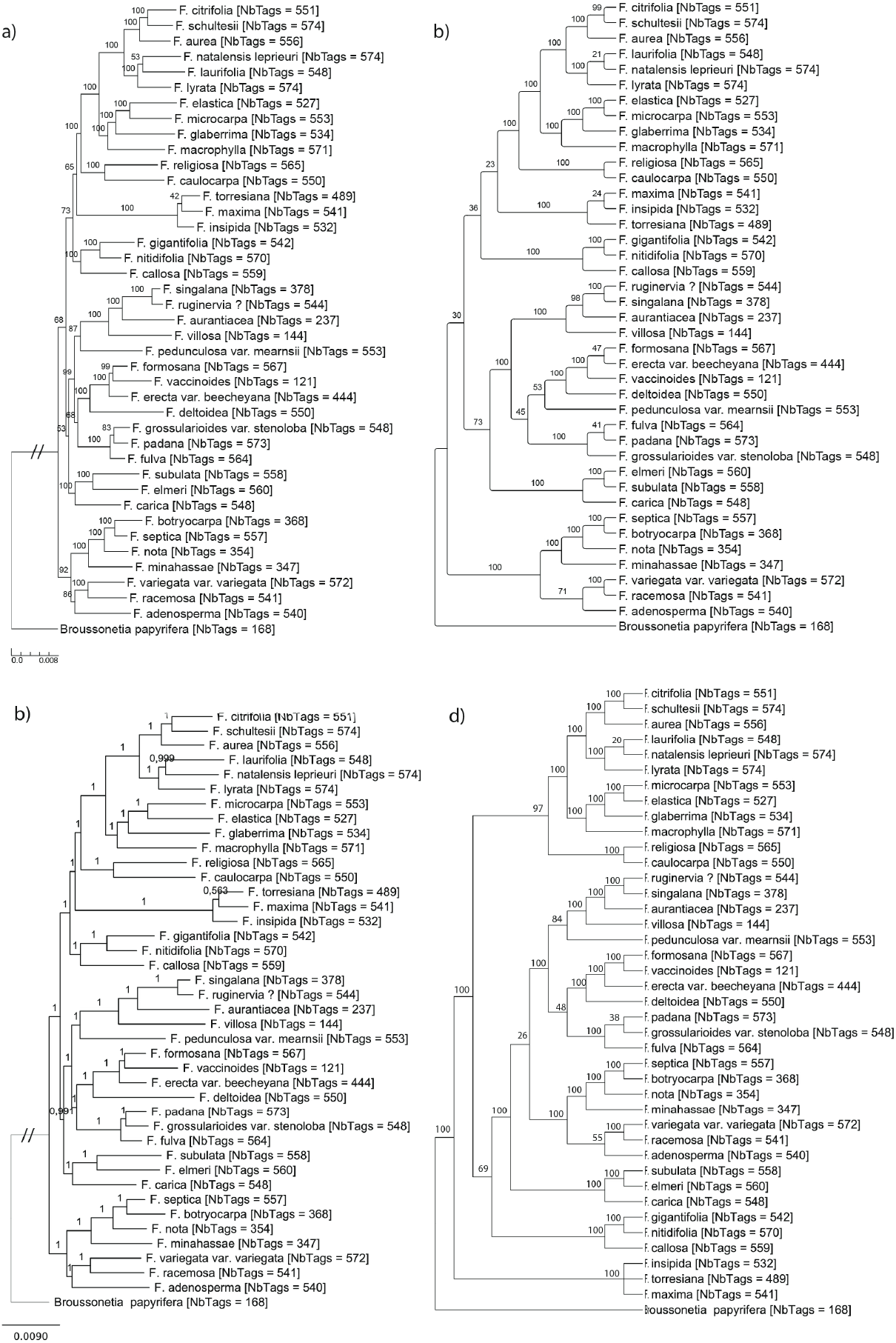
Phylogenetic trees obtained with the different analytical approaches. The number of RAD tags analysed for each sample is indicated between brackets. a) RAxML tree (Bootstrap values at nodes; 100 replicates); b) ASTRAL tree (Bootstrap values at nodes; 100 replicates); c) MrBayes tree (Posterior Probabilities at nodes); d) TNT tree (Bootstrap values at nodes; 100 replicates); strict consensus of the 2 most parsimonious trees, score = 25,089, CI = 0.725, RI = 0.683.

### Appendix S1: Supplementary methods

#### DNA extraction and RADseq library construction

Forty species of *Ficus* representing all known subgenera and 17 of the 20 sections as well as one outgroup were included in the study. It is noteworthy that the 3 missing sections (*Bosscheria, Dammaropsis, Papuasyce*) all belong to the subgenus *Sycomorus* which is undoubtedly monophyletic based on morphological characters. Leaves were sampled in the field, garden centres or own Herbarium by the authors or close collaborators. Leaves were either dried with silica gel or sun-dried.

20 mg of dried leaves were placed in Eppendorf vials and crushed with ceramic beads in liquid nitrogen. DNA was extracted with the Chemagic DNA Plant Kit (Perkin Elmer Chemagen, Baesweller, DE, Part # CMG-194), according to the manufacturer’s instructions with a modification of the cell lysis. The protocol was adapted to the use of the KingFisher Flex™ (Thermo Fisher Scientific, Waltham, MA, USA) automated DNA purification workstation. The powder was suspended in 400uL Lysis buffer (200mM Tris pH = 8.0, 50mM EDTA, 500mM NaCl, 1.25 % SDS, 0,5 % CTAB 1% PVP 40000, 1 g/100ml Sodium Bisulfite) and incubated 20 min at 65°C. Then 150 μL of cold precipitation buffer (sodium acetate 3M, pH 5.2) was added. Samples were centrifuged 10 min at 12000 rpm and 350 μL of the supernatant were transferred in a 96 deepwell plate. The binding of DNA on magnetic beads, the use of wash buffers and the elution of purified DNA were done following the Chemagic kit protocol and the use of the KingFisher Flex.

Library construction followed Baird et al. [1] and Etter et al. [2] with modifications detailed in Cruaud et al. [3] and below. An eight-cutter restriction enzyme was chosen (*SbfI*) to target conserved regions. The number of expected cut sites was estimated with the radcounter_v4.xls spread sheet available from the UK RAD Sequencing Wiki (www.wiki.ed.ac.uk/display/RADSequencing/Home). We assumed a 704 Mb approximate genome size ([4], ca 1.44 pg in average for 15 species of *Ficus*) and a 48% GC content (estimated from EST data available on NCBI). Based on those estimates, 9,095 cut sites were expected. This study was part of a larger project. Consequently, the library included more samples (128) than the 41 analysed here. 125ng of input DNA was used for each sample. After digestion, 1 uL of P1 adapters (100nM) were added to saturate restriction sites. Samples were then pooled sixteen by sixteen and DNA of each pool was sheared to a mean size of ca 400 bp using the Bioruptor^®^ Pico (Diagenode) (15sec ON / 90sec OFF for 8 cycles). After shearing, end repair and 3’-end adenylation, DNA of each pool was tagged with a different barcoded P2 adapter. A PCR enrichment step was performed prior to KAPA quantification. 2*125nt paired-end sequencing of the library was performed at MGX-Montpellier GenomiX on one lane of an Illumina HiSeq 2500 flow cell.

#### Phylogenetic inferences

Phylogenetic analyses were performed with maximum likelihood (RAxML [5]), Bayesian (MrBayes-3.2.6, [6]) and parsimony (TNT, [7]) approaches (concatenate, unpartitioned data sets) as well as gene tree reconciliation method (ASTRAL-III, [8]). For the RAxML analysis (raxmlHPC-PTHREADS-AVX version 8.2.4), a rapid bootstrap search (100 replicates) followed by a thorough ML search (-m GTRGAMMA) was performed. For Mrbayes analyses, model parameter values were initiated with default uniform priors and branch lengths were estimated using default exponential priors. A GTR + Gamma (4 categories) model was used. To improve mixing of the cold chain and avoid it converging on local optima, we used Metropolis-coupled Markov chain Monte Carlo (MCMCMC) simulation with each run including a cold chain and three incrementally heated chains. The heating parameter was set to 0.02 in order to allow swap frequencies from 20% to 70%. We ran two independent runs of 5 million generations with parameter values sampled every 500 generations. Convergence was assessed with Tracer 1.7 [9] and a consensus tree was built from the pooled samples from the stationary phases of the two independent runs (25% burnin). For the TNT analysis 100 searches were conducted that implemented random sectorial searches, 5 rounds of tree-drifting and tree-fusing and 50 ratchet iterations, holding up to 10 trees for each search until the best score was hit 50 times. Resulting trees were then subjected to TBR branch-swapping and a strict consensus was generated. Node support was computed with 100 bootstrap replicates. For the ASTRAL-III analysis, individual trees were inferred from each locus using RAxML (rapid bootstrap search 100 replicates and thorough ML search) and 100 multi-locus bootstrapping (MLBS, site-only resampling [10]) were conducted. Trees were annotated with TreeGraph 2.13 [11]. Summary statistics for data sets and samples were calculated using AMAS [12]. Analyses were performed on a Dell PowerEdge T630 with 10 Intel Xeon E5-2687 dual-core CPUs (3.1 GHz, 9.60 GT/s), 125 Go RAM and 13 To hard drive and on the Genotoul Cluster (INRA, Toulouse).

### Evolution of life history traits

Three key traits (plant habits, monoecy-gynodioecy and pollination syndromes) were studied. Stochastic mapping [13] as described in Bollback [14] and implemented in the R package phytools 0.6 [15] was utilized to estimate the ancestral state and the number of transitions for each trait. The fully resolved MCC tree obtained from the Bayesian analysis of the dataset was taken as input tree (outgroup pruned). The transition matrix was first sampled from its posterior probability distribution conditioned on the substitution model (10,000 generations of MCMC, sampling every 100 generations). Then, 100 stochastic character histories were simulated conditioned on each sampled value of the transition matrix. Three Markov models were tested: equal rates model (ER) with a single parameter for all transition rates; symmetric model (SYM) in which forward and reverse transition have the same rate and all rates different model (ARD). AIC scores and Akaike weight for each model were computed.

